# Extensive endemic transmission of multidrug-resistant *Mycobacterium tuberculosis* in Bhutan: A retrospective genomic-epidemiological study

**DOI:** 10.1101/2025.08.05.668595

**Authors:** Thinley Dorji, Karchung Tshering, Lila Adhikari, Thinley Jamtsho, Pavitra Bhujel, Pema Lhaden, Sonam Wangchuk, Wytamma Wirth, Kristy Horan, Justin T Denholm, Norelle L. Sherry, Michelle Sait, Timothy P. Stinear, Benjamin P. Howden, Patiyan Andersson

## Abstract

Despite decreasing overall tuberculosis notifications, the proportion of multidrug-resistant tuberculosis (MDR-TB) cases are increasing in Bhutan. While most MDR-TB cases are diagnosed among patients in the bordering districts and the capital, current diagnostic tests are limited in their ability to differentiate between the recurrent introductions and local transmission. For the first time, we conducted a retrospective genomic-epidemiological study to provide insights into the population structure, genotypic resistance patterns, and explore recent transmission of drug-resistant TB in Bhutan. Whole genome sequencing was performed on randomly selected drug-resistant and drug-sensitive TB isolates from Bhutan, collected between 2018-2022 at Microbiological Diagnostic Unit Public Health Laboratory in Melbourne, Australia. We investigated drug resistance mutations, and genomic clustering of cases using different single nucleotide polymorphisms (SNPs) thresholds. Of the 203 sequences that passed the quality control, 126 (62.1%) were MDR-TB and 15 (7.4%) were isoniazid-resistant TB. There were four different circulating lineages, with most sequences belonging to lineage 2 (86.2%). Using a SNP-threshold of ≤12 SNPs, 71% of sequences formed 12 genomic clusters. Surprisingly, the largest cluster included of 88% of all MDR-TB sequences and spanned the entire study period. These cases were highly clonal (mean pairwise SNP-distance of 10, range 0-25). Phylogenetic analysis with publicly available international sequence data showed that this MDR-TB cluster formed a distinct clade. The major burden of MDR-TB in Bhutan appears to be due to recent local transmission of cases resulting in a large, single endemic cluster. This information will be critical for TB control program in Bhutan to tackle the major burden of MDR-TB through enhanced contact tracing of this MDR-TB clade. Additionally, this genomic data will be valuable to regional neighbours to monitor for dissemination of the strain. This study highlights the significant value of investing in TB genomics in resource limited settings globally to gain actionable insights into transmission dynamics.

## Introduction

Timely and accurate diagnosis of tuberculosis (TB) is critical for effective treatment. However, these efforts are impeded by the slow growth of *Mycobacterium tuberculosis* (*Mtb)* and increasing cases of multidrug-resistant/rifampicin resistant TB (MDR-TB/RR-TB). While previous exposure and non-adherence to treatment were thought to be the major drivers for MDR-TB, current evidence suggests that majority of cases are now due to transmission of drug-resistant clones [1]. Therefore, the most impactful public health intervention will be disruption of MDR-TB transmission chains.

Whole genome sequencing (WGS) of *Mtb* has been used to describe the drug resistance and characterise the nature of transmission events [2]. It has high sensitivity for detection of *Mtb* clusters [3] and inferring antibiotic resistance patterns, with potential to detect more resistance than phenotypic drug susceptibility testing (pDST) or current WHO-endorsed genotypic tests [4]. Currently, WGS resistome inference has been endorsed for all first line TB drugs and some second line drugs, supported by a catalogue of mutations compiled by WHO [5].

Bhutan, a small landlocked country has a high burden of TB with an incidence rate of 164 per 100,000 population. It shares a border with India and China, where the incidence of TB was estimated to be around 195/100,000 population and 59/100,000, respectively in 2020 [6]. We have previously documented high incidence of MDR-TB in Bhutan, with 9% among new cases and 21% among previously treated cases [7]. While the country has seen decreasing overall notifications of TB, the proportions of MDR-TB has increased from 1.3% (17/1,349) in 2010 to 7.1% (65/919) in 2020 [8]. Epidemiological assessments show MDR-TB is spread across all districts of Bhutan, with most cases diagnosed among people living in the capital and southern districts bordering India [7]. To guide effective control strategies, it is essential to understand the prevailing resistance patterns and whether MDR-TB arises from multiple introductions or local transmission of a few adapted strains. We therefore conducted a retrospective genomic study of DR-TB cases from 2018–2022 to examine population structure and transmission dynamics.

## Methods

### Study settings

In Bhutan, TB samples are routinely collected from all presumptive TB cases in district or sub-district hospitals and undergo acid-fast microscopy and Xpert MTB/RIF testing (where available). Samples positive for either test, or smear negative highly suspicious clinical samples, are shipped to the National Tuberculosis Reference Laboratory (NTRL), Royal Centre for Disease Control. These samples undergo culture in mycobacterial growth indicator tubes (MGIT) and pDST for rifampicin, isoniazid, ethambutol and streptomycin.

Additionally, GenoType MTBDRplus is used to test for rifampicin and isoniazid resistance, and MTBDRsl for resistance to fluoroquinolone and amikacin/kanamycin. Patient details and sample data are recorded in the web-based Tuberculosis Information Surveillance System (TBISS).

### Sample selection

We sequenced samples received from the NTRL, that were identified to be DR-TB by pDST or MTBDRplus from 2018 to 2022. The proportion of the sequenced data, based on phenotypic test or rapid genotypic test performed at NTRL, against total drug-resistant samples present in NTRL is shown in Supplementary figure 1. Higer proportion of MDR-TB (range from 37% to 66% for a given year) and pre-XDR-TB (range from 33% to 100%) were available for sequencing. Additionally, some randomly selected drug-sensitive samples were also sequenced for the phylogenetic analysis.

### DNA extraction and sequencing methods

The sub-cultures were shipped from NTRL to the Microbiological Diagnostic Unit (MDU) Public Health Laboratory, Doherty Institute (Melbourne, Australia) in 7H9 liquid broth. DNA was extracted directly from the sub-cultures shipped from NTRL (without further culture) by bead-beating followed by sodium acetate precipitation of the DNA [9]. Library preparation was performed using Nextera XT preparation kit as per the manufacturer’s instructions and sequencing performed as previously described [10] on the Illumina NextSeq platform.

### Bioinformatic analysis

The quality checks and trimming of the FASTQ files were performed using the default settings of fastp [11] and species identified using Kraken-2 [12]. We used TB-profiler v.4.2.0 [13] for detection of resistance conferring mutations and determination of the lineages. Bohra (https://github.com/MDU-PHL/bohra), an in-house pipeline (certified to ISO standards in our laboratory) was used for isolate characterisation and phylogenetic analysis. Bohra uses snippy (https://github.com/tseemann/snippy) to make a core alignment to reference genome H37Rv (GenBank accession no. NC_000962). SNPs in repetitive regions were masked and sequences with alignment <95%, depth <30X, and mixed lineages were excluded from further phylogenetic analysis. Maximum likelihood phylogenetic tree was constructed using concatenated SNPs from multiple sequence alignment using IQTREE with constant sites with GTR + G4 and 1000 bootstraps [14]. To investigate geospatial patterns, we analysed publicly available 3,314 high-quality lineage 2 sequences from neighbouring countries and CRyPTIC study [15] (Supplementary table 1).

### Cluster analysis

Pairwise SNP-distance matrix was calculated using snp-dists (https://github.com/tseemann/snp-dists). As established in previous studies, we used single-linkage agglomerative hierarchical clustering with SNP distance threshold of ≤5 SNPs and ≤12 to define cases attributed to recent transmission [3, 16]. It is predicted that *Mtb* would acquire 5 SNPs in three years, and 10 SNPs in 10 years [3]. Isolates below this threshold were regarded as potentially belonging to a transmission cluster, although direction of transmission cannot be solely determined based on genomic data.

### Statistical analysis

All statistical analyses were performed using R 4.5.0. Packages used for data wrangling and visualisations include; dplyr [17], ggplot2 [18], ggtree [19], pheatmap [20], gtsummary [21], igraph [22], ape [23], and phangron [24]. The Fisher exact test was used to analyse the categorical data, while Wilcoxon-rank sum test was applied for continuous data. We performed multiple logistic regression using base R *glm* to calculate odds ratio and 95% confidence interval to identify variables associated with clustering. Model fitness was assessed with likelihood ratio test and backward stepwise regression [25]. Additionally, multicollinearity was tested using variance inflation factor (VIF) and all variables had VIF less than 5.

## Results

### Mycobacterium tuberculosis population structure

Of the 167 DR-TB and 53 DS-TB isolates sequenced (Figure 1), 14 sequences with contamination, one duplicate sequence, and two samples with mixed *Mtb* lineages were excluded from the subsequent analysis. This resulted in a final dataset of 203 *Mtb* sequences, with 92% of sequences passing our quality control. This was despite sequencing directly from the sub-cultures shipped from NTRL. Reads were mapped to the H37Rv reference genome, with mean read coverage of 96.6% and average read depth of 154 (Range: 36.4 - 275).

**Figure 1.**
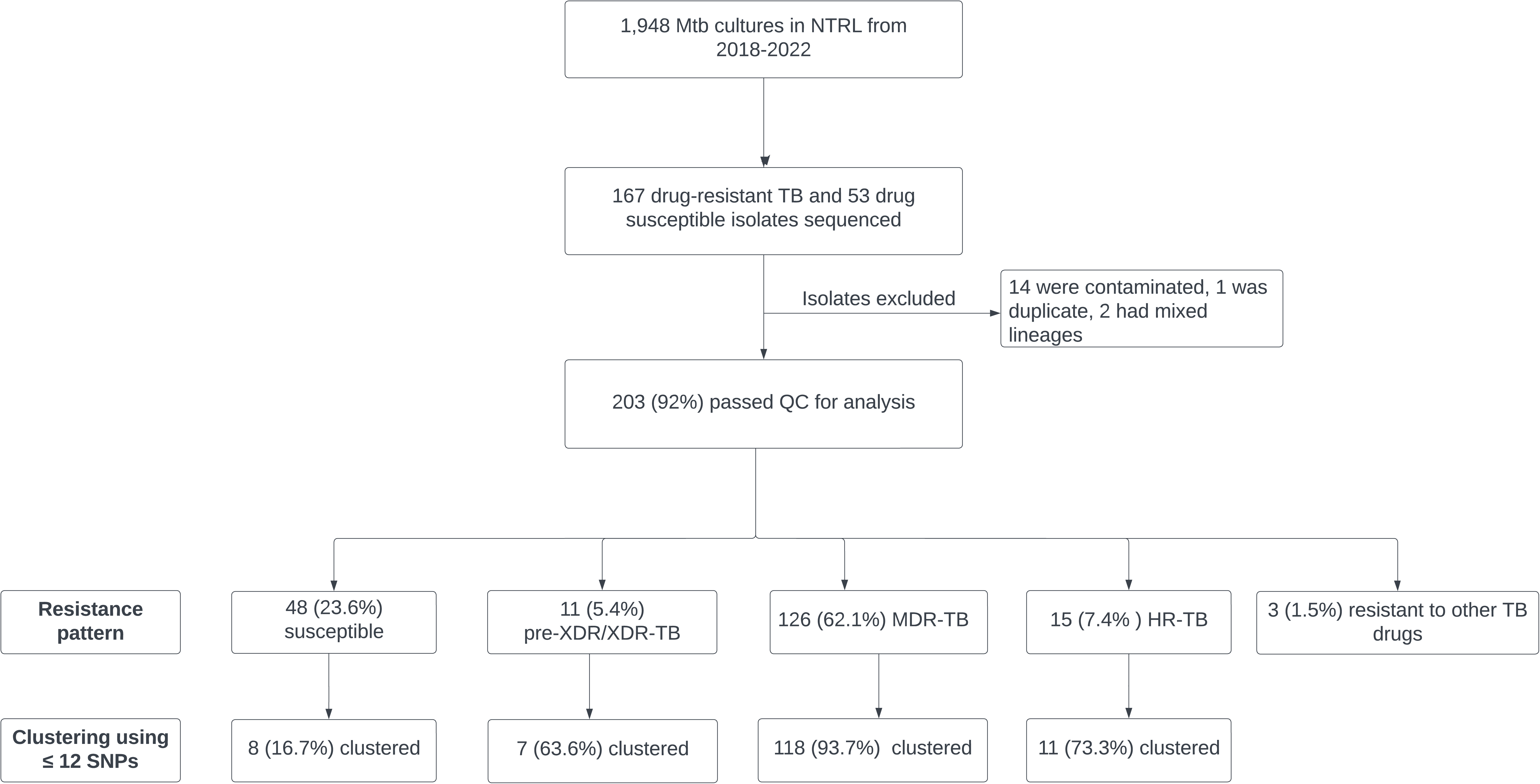
Flow diagram of samples tested, and resistance patterns observed. The 167 drug-resistant and 53 drug-sensitive were selected based on phenotypic drug susceptibility test or genotypic line probe assay. MDR-TB (-drug resistant TB), HR-TB (isoniazid resistant TB), pre-XDR/XDR-TB (pre-extensively drug-resistant TB/ extensively drug-resistant TB). Others include streptomycin resistant TB and fluoroquinolones resistant TB. NTRL (National Tuberculosis Reference Laboratory).

Associated epidemiological data indicated the samples were representative of most districts of Bhutan (Supplementary figure 2).

### Demographic characteristics

Female cases accounted for higher proportion of our sequenced samples (n=115, 56.7%). The median age of patients was 26 years (range 6-86 years). The majority of our sequenced samples (n=183, 90.1%) were pulmonary bacteriologically confirmed TB. While 76.8% (n=156) of the cases were new TB cases, 14.8% (n=30) were previously treated for TB. Most cases were students (n=54, 26.6%) followed by business (n=29, 14.3%).

### Genomic diversity of *M. tuberculosis* in Bhutan

A maximum likelihood phylogenetic tree was inferred using a core genome alignment of the 203 isolates, with 8,006 high confidence SNPs. The majority of sequences belonged to *Mtb* lineage 2 (East Asian lineage, n=175 (86.2%)), which were all assigned to the sub-lineage 2.2.1 (Figure 2). This lineage had Thr2Ala *esxW* mutation, which is associated with increased transmissibility [26]. This was followed by lineage 4 (Euro-American lineage, n=16 (7.9%)) and had highest diversity among all lineages. There were small numbers of lineage 3 (East African Indian lineage, n=7 (3.4%)) and lineage 1 (Indo-Oceanic lineage, n=5 (2.5%)).

**Figure 2.**
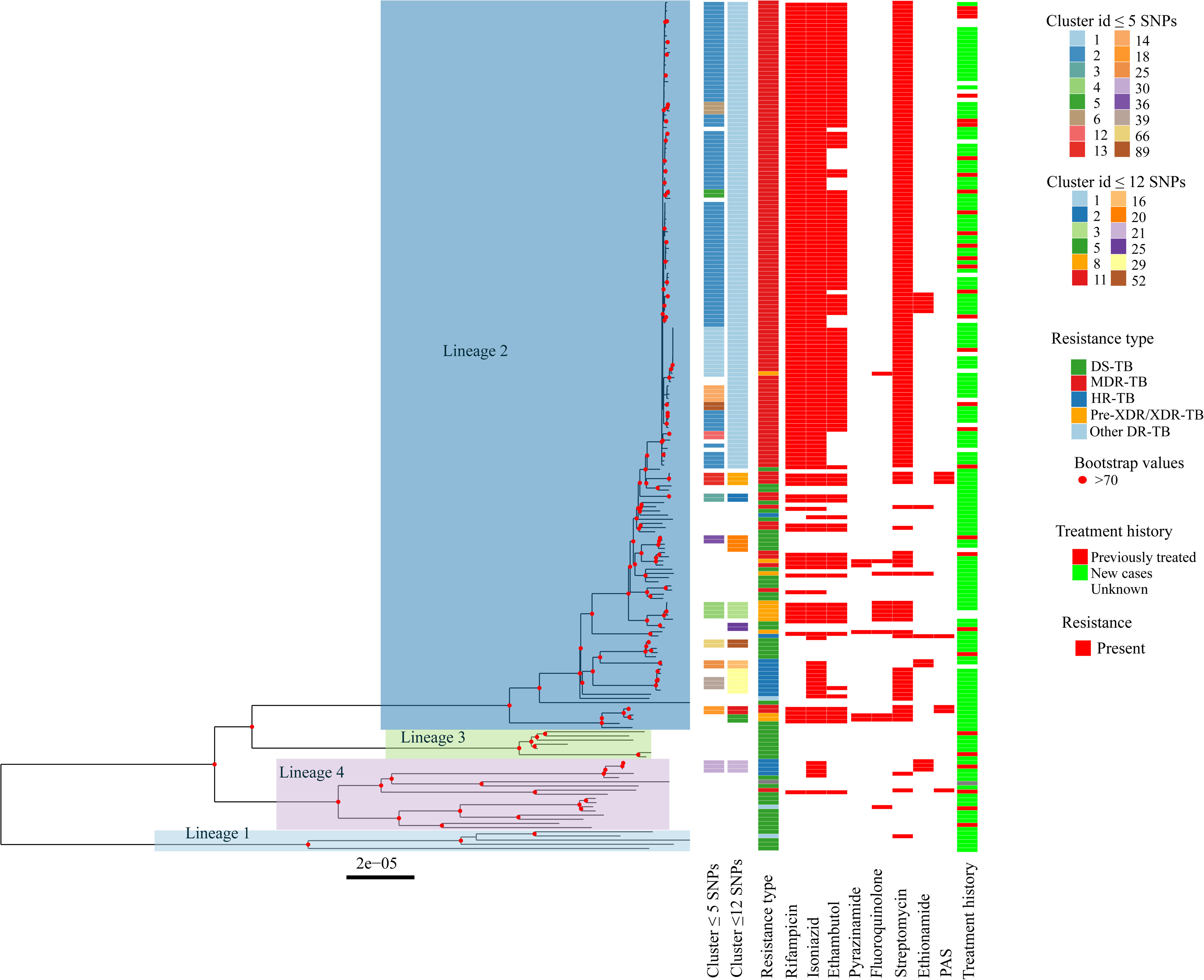
Midpoint rooted maximum likelihood phylogenetic tree of 203 *Mycobacterium tuberculosis* sequences. The tree is highlighted with lineages. From left to right, the heatmap shows clustering using SNPs thresholds of ≤5, ≤12 and resistance type. Rest of the heatmap shows the presence or absence of resistance. The last heatmap shows the treatment history. DS-TB (drug sensitive TB), MDR-TB (multidrug-resistant TB), HR-TB (isoniazid resistant TB), pre-XDR/XDR-TB (pre-extensively drug-resistant TB/ extensively drug-resistant TB), other DR-TB (drug-resistant TB) includes resistance to other TB drugs than rifampicin and isoniazid, PAS (para-aminosalicylic acid).

While WGS is validated for detection of resistance to first line TB drugs and some second line drugs[27], it is not validated for group A drugs such as bedaquiline and linezolid.

Therefore, in this work pre-extensively drug-resistant TB (pre-XDR TB, a case of MDR-TB with additional resistance to fluoroquinolone) was grouped together with extensively drug-resistant TB (XDR-TB, a case of pre-XDR-TB and resistance to group A drugs) as pre-XDR/XDR-TB. Most sequences in our dataset were genomically identified as MDR-TB (n=126, 62.1%), isoniazid resistant TB (HR-TB, n=15, 7.4%) and pre-XDR/XDR-TB (n=11, 5.4%) (Figure 1). Among the MDR-TB sequences, 99.2% (n = 125) belonged to lineage 2.

While lineage 2 was significantly associated with MDR/pre-XDR/XDR-TB (OR = 91.97, 95% CI: 14.3-3794.4, p < 0.001), HR-TB was more likely to be associated with lineage 4 (OR = 5.26, 95% CI: 1.06-21.57, p = 0.02). All lineage 3 isolates were found exclusively among genomically sensitive *Mtb* sequences.

### Characterisation of *M. tuberculosis* antibiotic resistance mutations

Most isoniazid resistance mutations were seen in *katG* Ser315Thr (95.4%, n=145), which confers high level isoniazid resistance [28]. Five sequences had *inhA* c.-777C>T mutation (previously known as *fabG1* c.-15C>T), known to cause low to moderate level isoniazid resistance [4, 28] and was exclusively seen among HR-TB in our dataset. One MDR-TB sequence had dual *katG* Thr251Met and *inhA* c.-777C>T, with former being relatively uncommon and linked to moderate level resistance [29]. *katG* mutations, which confers high level isoniazid resistance is not typically associated with *inhA* promotor mutations and this co-occurrence is likely due to exposure to ethionamide [30].

All rifampicin resistance mutations in our dataset occurred in the 81-bp rifampicin resistance determining region (RRDR) (426-452) of *rpoB,* and any non-synonymous mutation in this region is presumed resistant [5]. Within this region, we identified 133 (97.1%) sequences with Ser450Leu, associated with high-level resistance [28]. Others include His445Tyr (n=1), and one sequence with dual Asp435Gly and Leu452Pro. The presence of a second *rpoB* mutation is known to increase the rifampicin MIC, leading to stepwise increase of resistance [28]. Leu452Pro mutations result in borderline rifampicin resistance [5] and are often missed during MGIT testing [31]. Only five different *rpoB* variants were seen in our sequences, possibly due to limited genetic diversity within the country, and intolerance of *rpoB* to major changes [32].

Up to 81.7% (n=103) of MDR-TB cases exhibited concurrent *embB* mutations with majority of them being Asp328Gly (n=30), Gly406Ala (n=24), Met306Val (n=22) (Supplementary figure 3). While Gly406Ala and the Met306Val are associated with ethambutol resistance, Asp328Gly is classified as having “uncertain significance” [5], according to the WHO catalogue of mutations. Fluoroquinolone resistance conferring mutations were identified in 6% (n =12) of the sequences, all of which were located within the quinolone resistance-determining region (QRDR) of *gyrA* (codons 74-113). The most common mutation among them was Asp94Tyr (n = 7). All pre-XDR/XDR-TB sequences were also resistant to streptomycin and ethambutol.

### Inference of potential *Mtb* transmission clusters

The overall median pairwise SNP-distance among our 203 sequences was 90 SNPs (Range: 0-1700). Lineage 2 sequences had lowest median SNP-distance of 66 SNPs followed by lineage 3 with 217 SNPs, lineage 1 with 523 SNPs and lineage 4 with 628 SNPs (Supplementary figure 4).

Using a SNP threshold of ≤5 SNPs, 63.5% (129/203) of sequences were grouped into 16 clusters, suggesting recent transmission within last three years [3]. However, when we increased the threshold to ≤12 SNP, the clustering rate increased to 70.9% (144/203) with formation of 12 clusters. At this threshold, four clusters were formed by patients with MDR-TB (n=118/126; 93.7%), three clusters by patients with HR-TB (n=11/15; 73.3%) and three clusters by patients with drug-sensitive TB (n=8/48; 16.7%). There were two clusters of pre-XDR/XDR-TB (6/11; 54.5%) (Figure 3). Unexpectedly, the largest cluster (cluster 1) comprised of 111 MDR-TB genomes and one pre-XDR-TB genome, constituting 88% of all MDR-TB genomes and 54.7% of our total sequences. The median pairwise SNP distance in this largest cluster was 9 SNPs (Range: 0-25).

**Figure 3.**
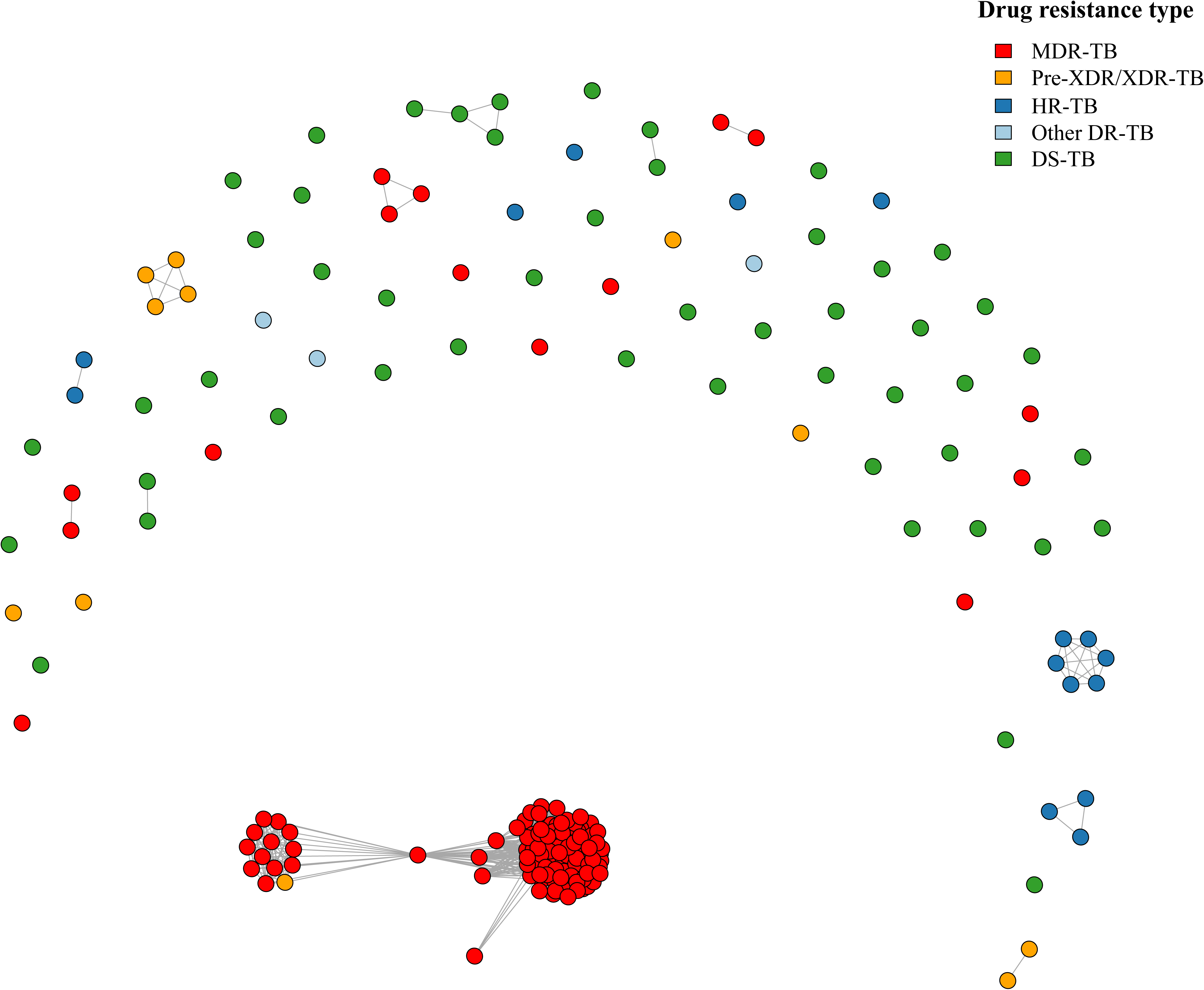
Clustering of cases using ≤12-SNPs threshold. Each dot represents a single sequence and coloured by resistance type. MDR-TB (multidrug-resistant TB), HR-TB (isoniazid resistant TB), pre-XDR/XDR-TB (pre-extensively drug-resistant TB/ extensively drug-resistant TB), DS-TB (drug sensitive TB), other DR-TB (drug-resistant TB) includes resistance to other TB drugs than rifampicin and isoniazid.

All clusters were from lineage 2, except for cluster 21, which belonged to lineage 4. This cluster was composed of three HR-TB sequences-two from a patient who was previously treated for TB and had recurrence after a year. The pairwise SNP distance between these two sequences was one SNP, indicating that this patient was not fully treated in the past and hence experienced relapse of TB.

Analysis of the spatial and temporal distribution of cases showed that most clusters were not confined to any specific geographical location, with multiple districts featuring in clusters (Figure 4A). The exceptions to this pattern were clusters 21 and 52. MDR-TB cluster 1 had the largest geographical distribution of cases (Figure 4B) throughout the country and spanned the entire study period, suggesting that transmission of this clade has been ongoing for some time in Bhutan. The most recent common ancestor (MRCA) of this cluster harboured *rpoB* Ser450Leu, *katG* Ser315Thr, and *rpsL* Lys43Arg, conferring resistance to rifampicin, isoniazid and streptomycin. Additionally, all isolates in this largest cluster carried the compensatory *rpoC* Leu516Pro mutation. This largest cluster was further sub-divided into seven smaller clusters using ≤5 SNP threshold, with some sub-clusters harbouring ethambutol resistance mutations (Supplementary figure 5). We attempted to explore the time of introduction and spread of the largest MDR-TB in Bhutan using Bayesian Evolutionary Analysis Sampling Trees (BEAST). However, this was not possible due to the absence of a temporal signal using probabilistic methods like Bayesian evaluation of temporal signal and clockor2 [33], likely due to slow mutation rate of *Mtb* and insufficient accumulation of mutations during study period.

**Figure 4.**
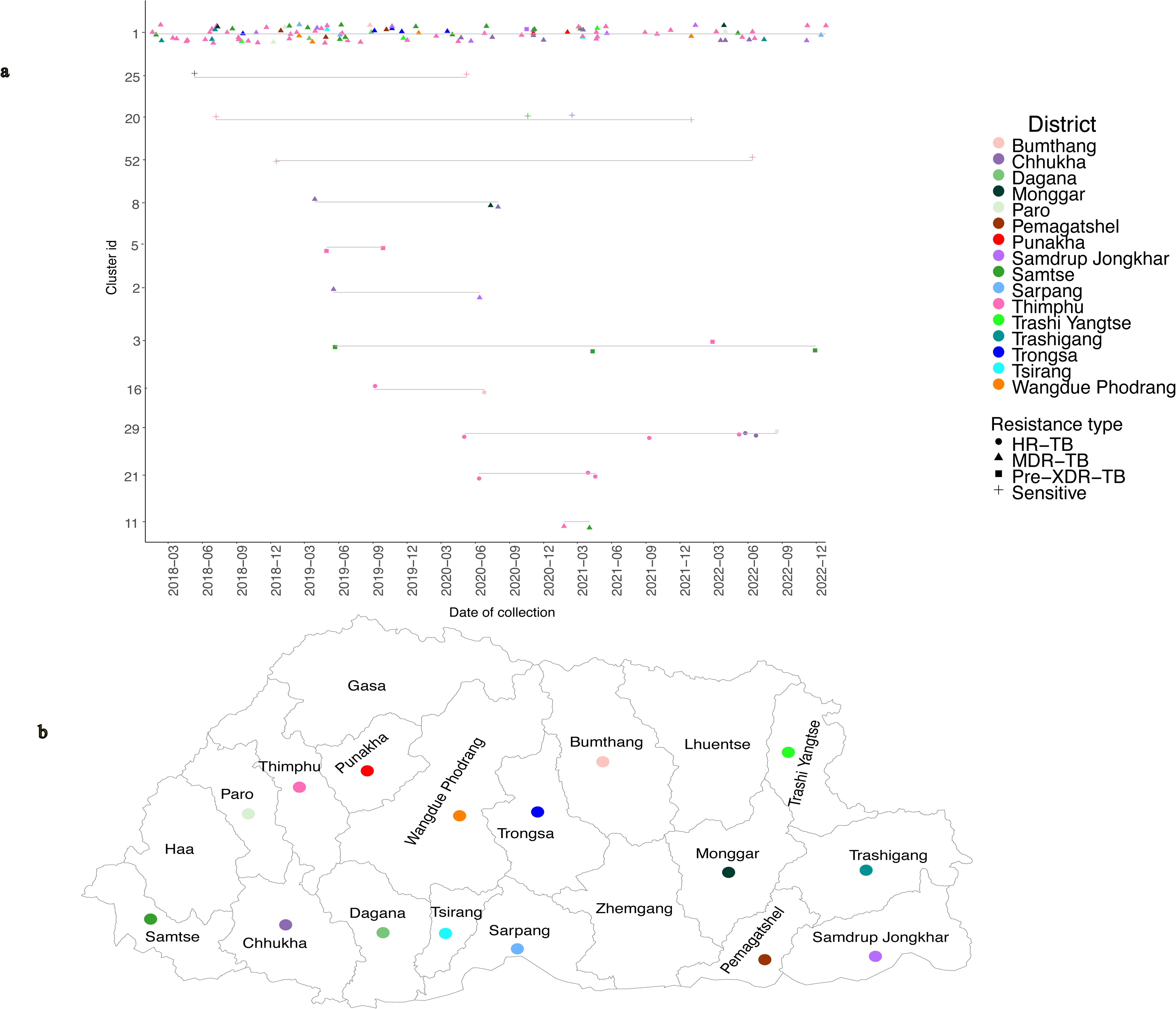
Distribution and clustering of TB cases by location of patients. (**a**) Cluster timeline of all clustered sequences based on the time of diagnosis and different cluster IDs. Each sequence is shown by different shapes indicating type of resistance pattern. Line shows that they belong to same cluster. The x-axis shows the date of collection of isolates. (**B)** The map shows the location of patients in the largest cluster. MDR-TB (multidrug-resistant TB), HR-TB (isoniazid resistant TB), pre-XDR/XDR-TB (pre-extensively drug-resistant TB/ extensively drug-resistant TB).

In our dataset, students represented the largest occupational group (n=54, 26.6%), with the median of 20 years (range: 14-28 years). We expected clustering of cases among students due to the potential for transmission in enclosed classrooms and dormitories. However, the clusters appeared quite heterogenous in terms of occupational groups (Supplementary figure 6), suggestive of community wide transmission.

### Factors associated with clustering

We investigated the demographic and bacterial factors that could likely lead to clustering of TB cases in Bhutan (Table 1). Clustering was more likely to seen among patients from Western region, possibly associated with high case numbers in this region. We also demonstrate significant clustering of TB cases among those diagnosed in years 2020 and 2021. This was likely influenced by COVID-19 pandemic. Lineage 2 sequences had higher odds of clustering compared to other TB lineages. Additionally, MDR-TB, pre-XDR/XDR-TB and HR-TB were also more likely to cluster, possibly influenced by a biased selection of predominantly DR-TB samples in our dataset. These data should be interpreted with caution as confidence intervals were wide, and further assessment with other epidemiologic data is required.

**Table 1.**
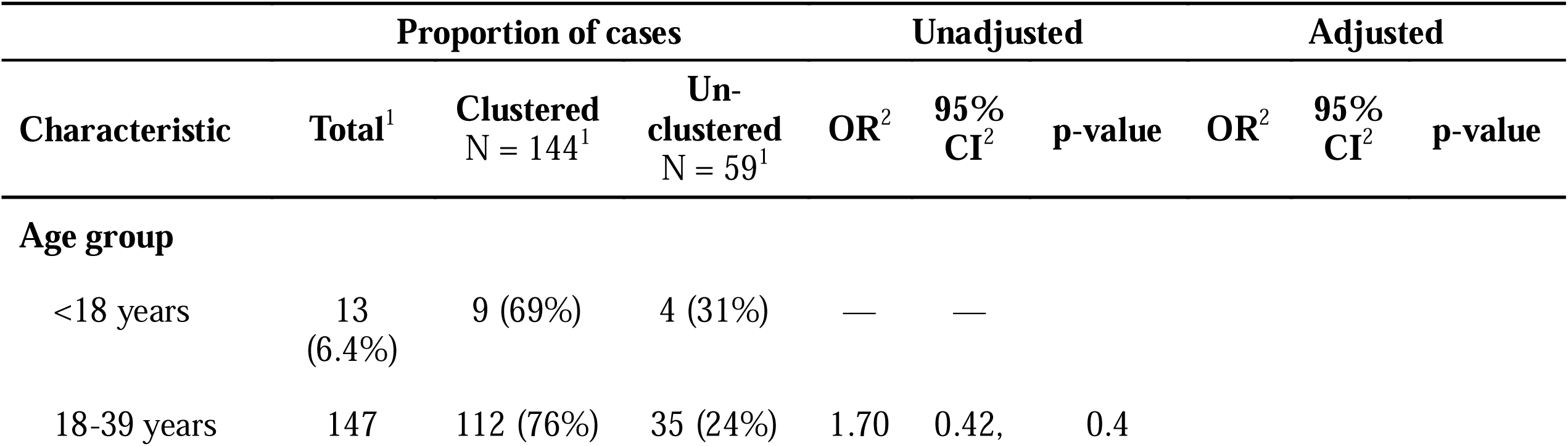
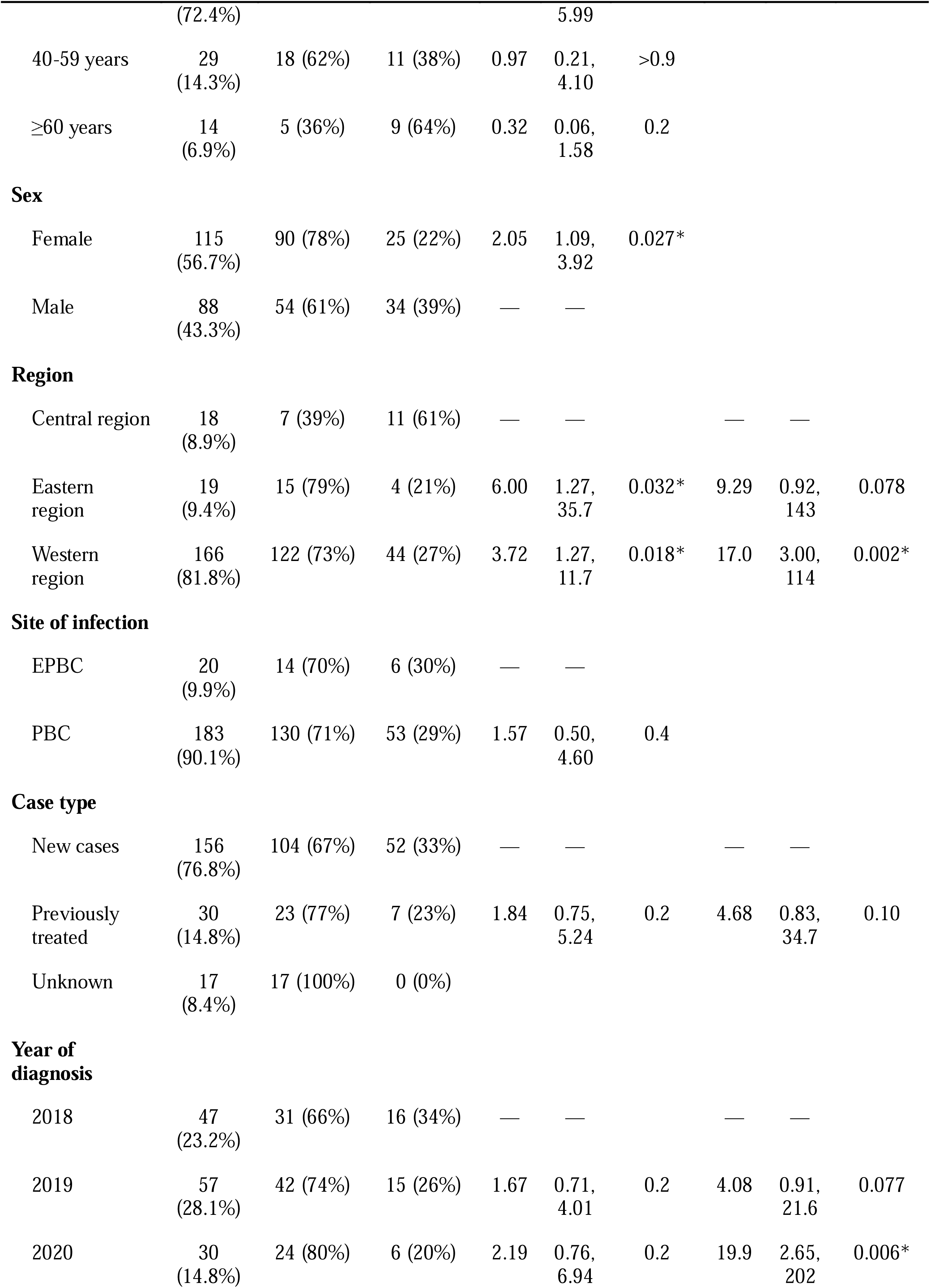
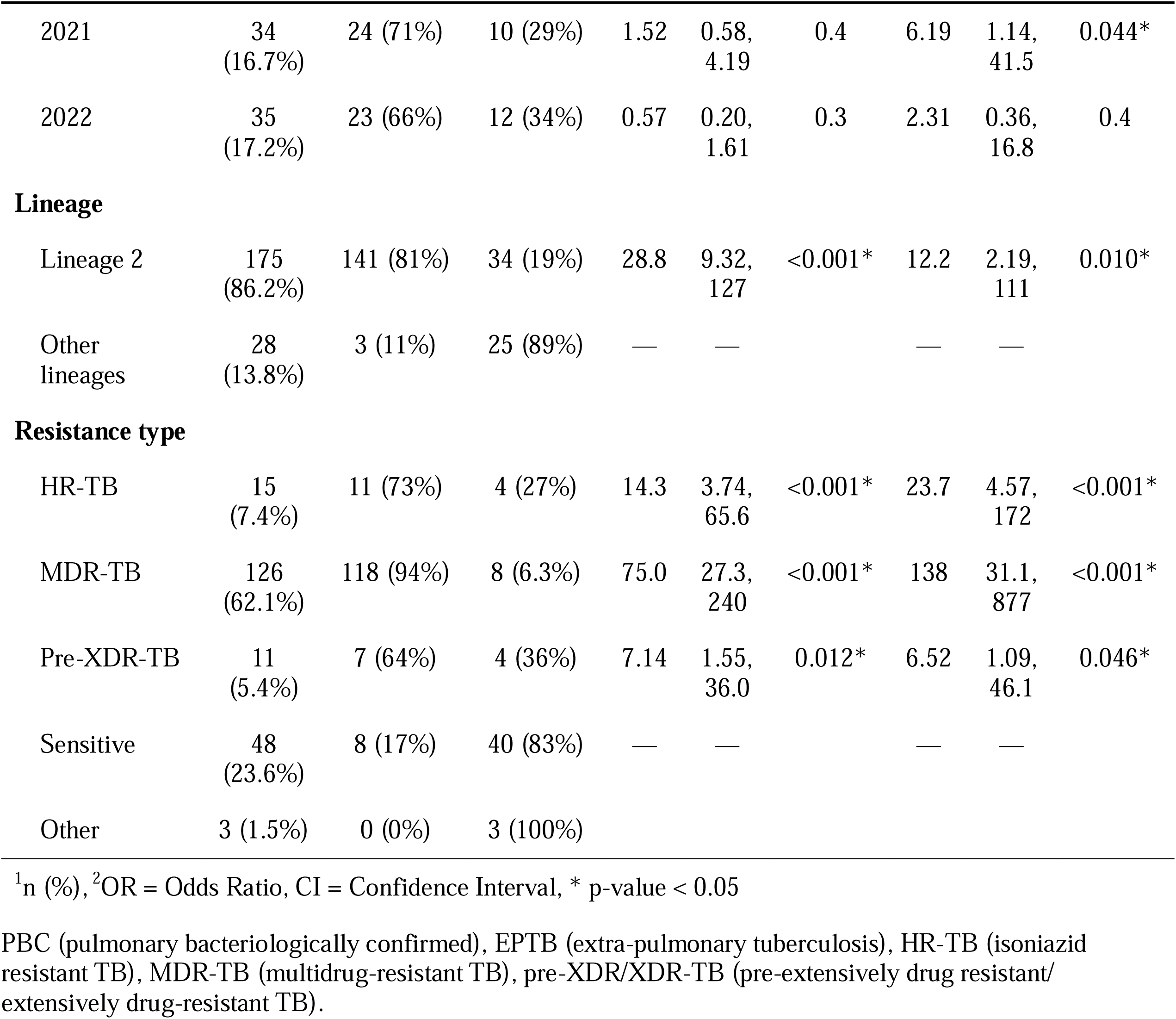
Factors associated with clustering of *Mtb* sequences.

### Contextualisation of Bhutan isolates to regional/global sequences

Given the predominance of lineage 2 in our sequences, we analysed the phylogenetic relationships with the publicly available lineage 2 datasets. Sequences from the largest Bhutanese cluster remained a monophyletic clade, consistent with a single introduction and recent expansion of this sub-lineage in Bhutan (Supplementary figure 7). The most closely related sequence was an MDR-TB from Bangladesh, with a pairwise SNP-distance of 23 from the nearest Bhutanese sequence.

## Discussion

To our knowledge, this is the first study investigating TB epidemiology in Bhutan using pathogen genomics. Remarkably, the major burden of MDR-TB is due to local endemic transmission, with a single large cluster accounting for 88% of all MDR-TB cases in this study. This cluster was not restricted to any geographical location or occupational group and spanned the entire study period. In the context of global TB, this MDR-TB cluster formed a separate Bhutan specific clade, suggesting its establishment and ongoing transmission among the local population. This finding marks a considerable shift in understanding of MDR-TB transmission in Bhutan with implications for public health intervention approaches.

Our dataset had predominance of sub-lineage 2.2.1, which is the dominant circulating sub- lineage in the Southeast Asia and has begun to replace other TB lineages [26]. This sub- lineage is associated with increased transmissibility, attributed to the presence of *esxW* mutation [26]. When we contextualised our lineage 2 sequences with global lineage 2 data, our largest cluster formed a distinct clade, with an MDR-TB strain from Bangladesh. While importation from Bangladesh cannot be ruled out, it is also possible that transmission originated from neighbouring Indian states, as Bhutan shares a long and open international border with India. Furthermore, Indian states bordering Bhutan also share borders with Bangladesh, raising the possibility that this MDR-TB strain could be circulating in the region. Unfortunately, at the time of study there were no publicly available sequences from the Indian states adjacent to Bhutan for comparison.

The clustering rate of 93.7% observed among our MDR-TB sequences is higher than that observed in studies from China [16] and India [34]. Our largest MDR-TB cluster spanned the entire study period, providing the evidence of sustained local MDR-TB transmission. Several factors indicate that this transmission has been ongoing for some time. Firstly, our commonest mutation *katG*-Ser315Thr and *rpoB*-Ser450Leu has been described among MDR-TB cases at high proportions since 2014 in Bhutan (77.9%) [35]. These mutations are associated with low fitness costs [32], and presence of the compensatory *rpoC* Leu516Pro [36] mutation in the largest MDR-TB cluster likely contributes to its successful spread.

Secondly, using 5-SNP threshold, the largest MDR-TB cluster was fragmented into multiple sub-clusters with diverse ethambutol mutations. This could be due to local evolution and adaptation with subsequent community transmission. The emergence of this strain likely resulted from MDR-TB misdiagnosis prior to the introduction of Xpert MTB/RIF in 2016, and the use of a drug-sensitive TB regimen to treat MDR-TB cases, inadvertently selecting for ethambutol resistance. Lastly, the largest cluster was seen among all occupational groups and was not localised to any specific district in the country.

As our study was retrospective, one limitation is the potential for selection bias in the isolates included in the analysis. Insights into the transmission dynamics of drug-sensitive TB samples were limited, due to the relatively small number of samples included. Routine prospective collection of *Mtb* isolates and genome sequencing is likely to impact clustering observed in the future. Our cluster analysis here is based on the sequencing data only, as the contact tracing data was not available. Thus, we cannot comment on transmission settings. In a future scenario of genomics-informed surveillance, such epidemiological information will be essential for contextualising genomic data and guiding optimal public health responses.

Despite these limitations, our findings provide compelling evidence that challenges previous assumptions and offers new directions for TB control efforts in Bhutan.

Previous studies demonstrated high incidence of MDR-TB cases in the bordering districts and the Capital [7] leading to the assumption that most MDR-TB cases were probably due to repeated cross-border transmissions into the country [37]. However, our finding refutes this hypothesis and provides evidence of local transmission of MDR-TB strain. Therefore, strengthening systematic contact tracing, screening for latent TB among contacts, and providing prophylaxis with fluoroquinolones [38] is an important strategy. This is further supported by the observation of low levels of fluoroquinolone resistance in our dataset. While screening of high-risk groups is cost-effective, population wide screening approaches can be considered, as they have the potential to significantly reduce TB incidence [39]. In light of the encouraging reductions in overall TB incidence, appropriately targeted public health strategies for tackling the increasing proportion of MDR-TB will continue the progress toward achieving the goals of TB elimination in Bhutan. More broadly, this study highlights the significant value of investing in TB genomics in resource limited settings globally to gain actionable insights into transmission dynamics.

## Ethical

The study was approved by the Research and Ethics Board of Health, Ministry of Health, Bhutan (*REBH/Approval/2022/027)* and University of Melbourne (Ethics approval *2023- 26130-41858-3*). The requirement for informed consent was waived due to the retrospective design of study, the use of deidentified data and deidentified bacterial isolates from patient specimens.

## Author contributions

TD, KH, BH and PA conceived and designed the study, and analysed the data. KT, LA, TJ, PB, PL and SW processed the samples, collected data and contributed to analysis of data. JD, NS, MS, WW and TS were involved in study design and analysis of the data. MS performed sequencing of samples and QC. All authors contributed to interpretation, drafting and approved the final manuscript.

## Supporting information

Publicly available sequences

## Acknowledgement

We are grateful to the National Tuberculosis Reference Laboratory, Royal Centre for Disease Control, Ministry of Health for their generous collaboration. We also acknowledge the efforts of the molecular laboratory team at Microbiological Diagnostic Unit Public Health Laboratory, Peter Doherty Institute of Infections and Immunity for sequencing.

## Supplementary figures

Supplementary figure 1. Proportion of different drug-resistant TB samples selected for sequencing by total of specific drug resistance samples per year available in National Tuberculosis Reference Laboratory based on MTBDRplus or phenotypic drug susceptibility test.

Supplementary figure 2. Distribution of lineages based on patient location. The number denotes the number of isolates sequenced from that district.

Supplementary figure 3. Upset plot showing the different combinations of resistance. The mutations are coloured based on the drug resistance and drug.

Supplementary figure 4. Comparison of pairwise SNP distances by lineage. The statistical difference in pairwise SNP-distance between lineages was calculated using pairwise Wilcoxon tests with Bonferroni correction.

Supplementary figure 5. Phylogenetic tree with resistance conferring mutations. The presence of mutations is coloured by red and absence with white. The resistance mutations and drugs are coloured same.

Supplementary figure 6. Clustering of the cases by occupation. This clustering was based on ≤12-SNPs threshold.

Supplementary figure 7. Comparison of Bhutanese TB sequences with global sequences. The tips and branches are coloured by the country of origin of sequences. The largest Bhutanese cluster is zoomed out to indicate the most common recent ancestor.

## Supplementary tables

Supplementary Table 1: Publicly available sequences used in this study

## Data availability

The genomic data from Bhutan used in this study are available in GenBank under accession PRJNA1241451. The sequences from other studies used in this study are listed in supplementary table 1.

