## Supplementary figures for "Extensive endemic transmission of multidrug-resistant *Mycobacterium tuberculosis* in Bhutan: A retrospective genomic-epidemiological study"


| Content | Page |
|---|---|
| **Supplementary figure 1.** Proportion of different drug-resistant TB samples selected for sequencing by total of specific drug resistance samples per year. | **1** |
| **Supplementary figure 2**. Distribution of lineages based on patient location. The number denotes the number of isolates sequenced from that particular district. | **2** |
| **Supplementary figure 3**. Upset plot showing the different combinations of resistance. | **3** |
| **Supplementary figure 4**. Comparison of pairwise SNP distances by lineage using pairwise Wilcoxon tests with Bonferroni correction. | **4** |
| **Supplementary figure 5**. Phylogenetic tree with resistance conferring mutations. | **5** |
| **Supplementary figure 6**. Clustering of the cases by occupation based on ≤12-SNPs threshold. | **6** |
| **Supplementary figure 7**. Comparison of Bhutanese TB sequences with global sequences. | **7** |

### Supplementary figure 1

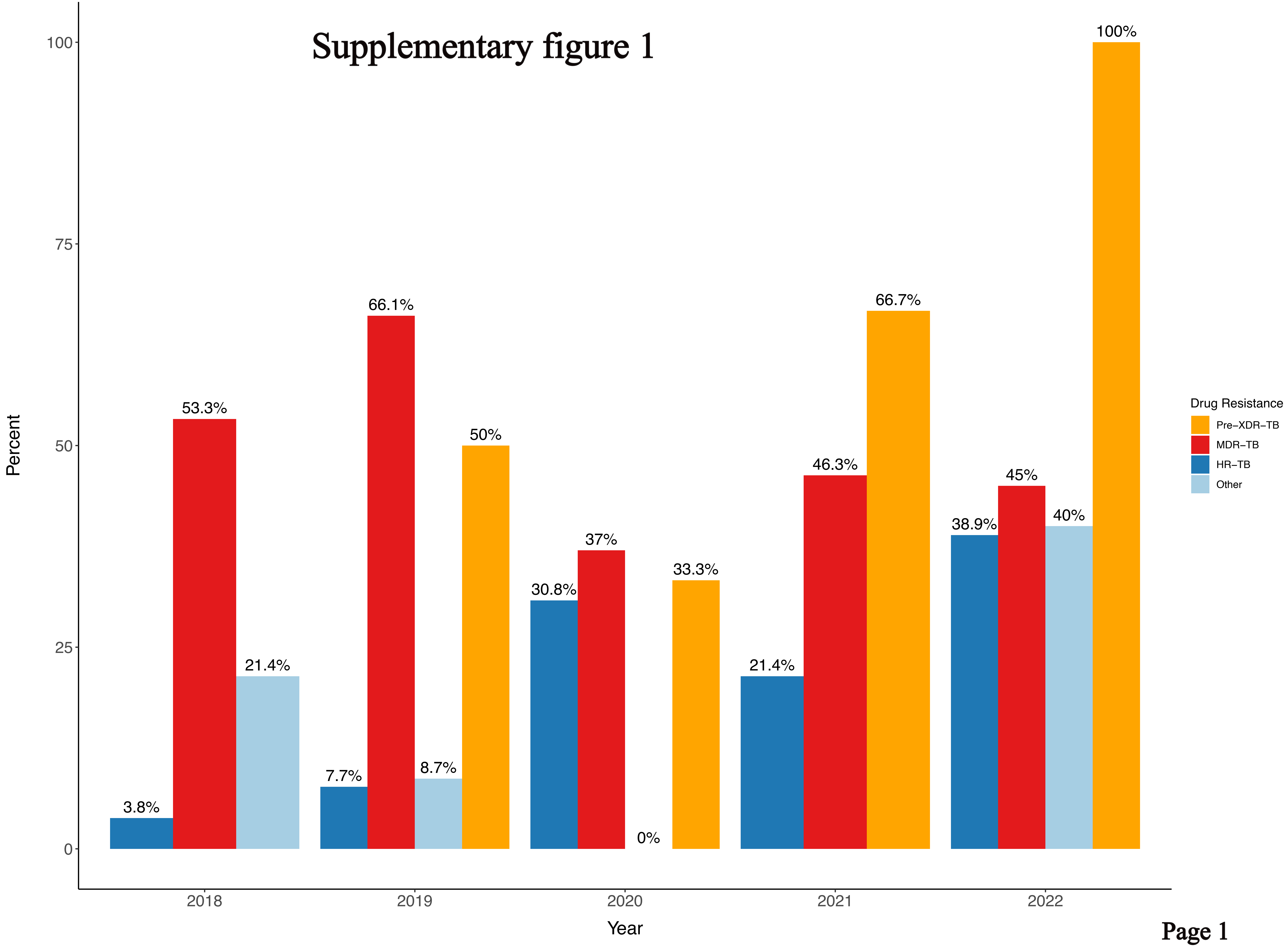

### Supplementary figure 2

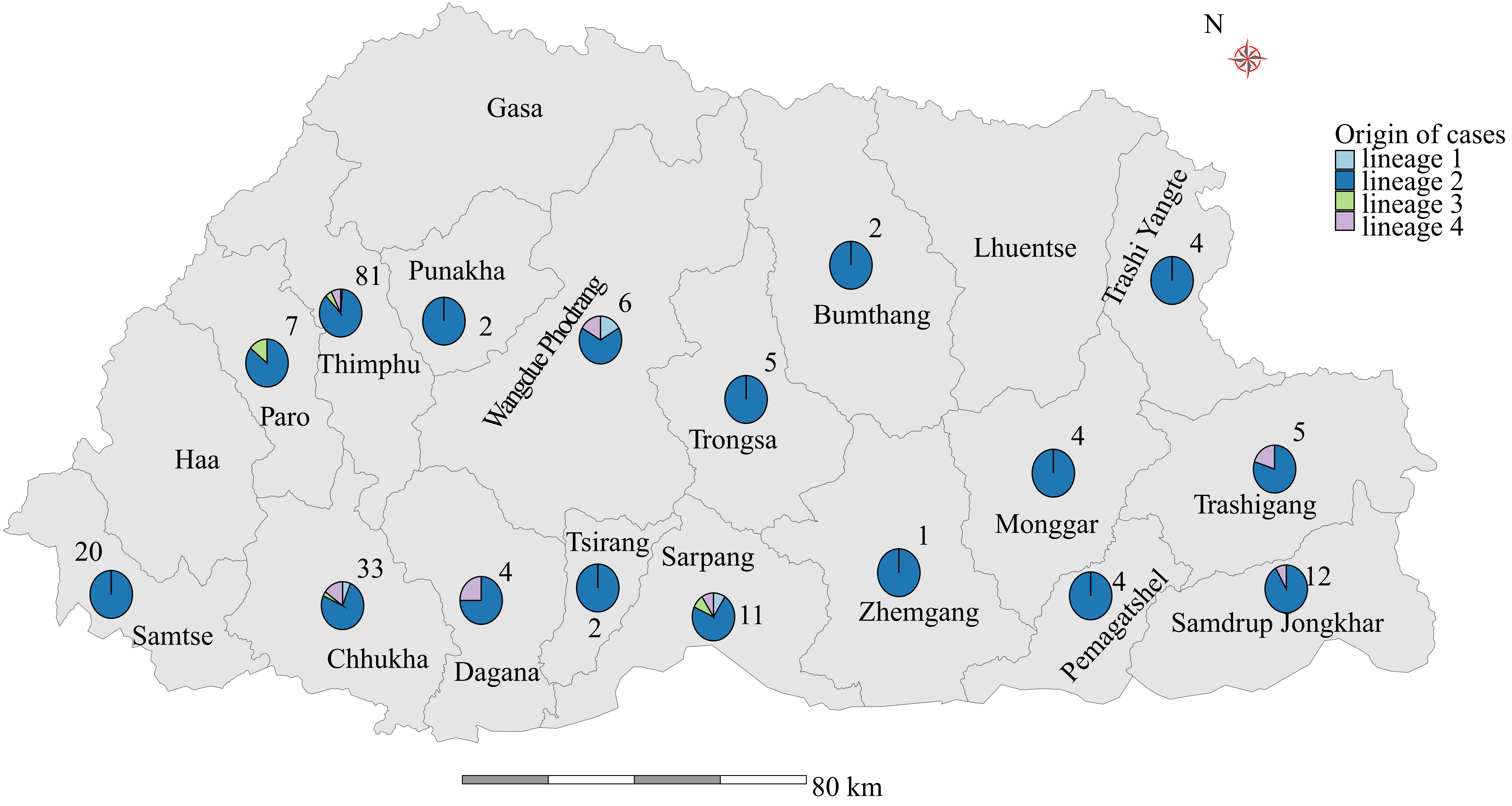

### Supplementary figure 3

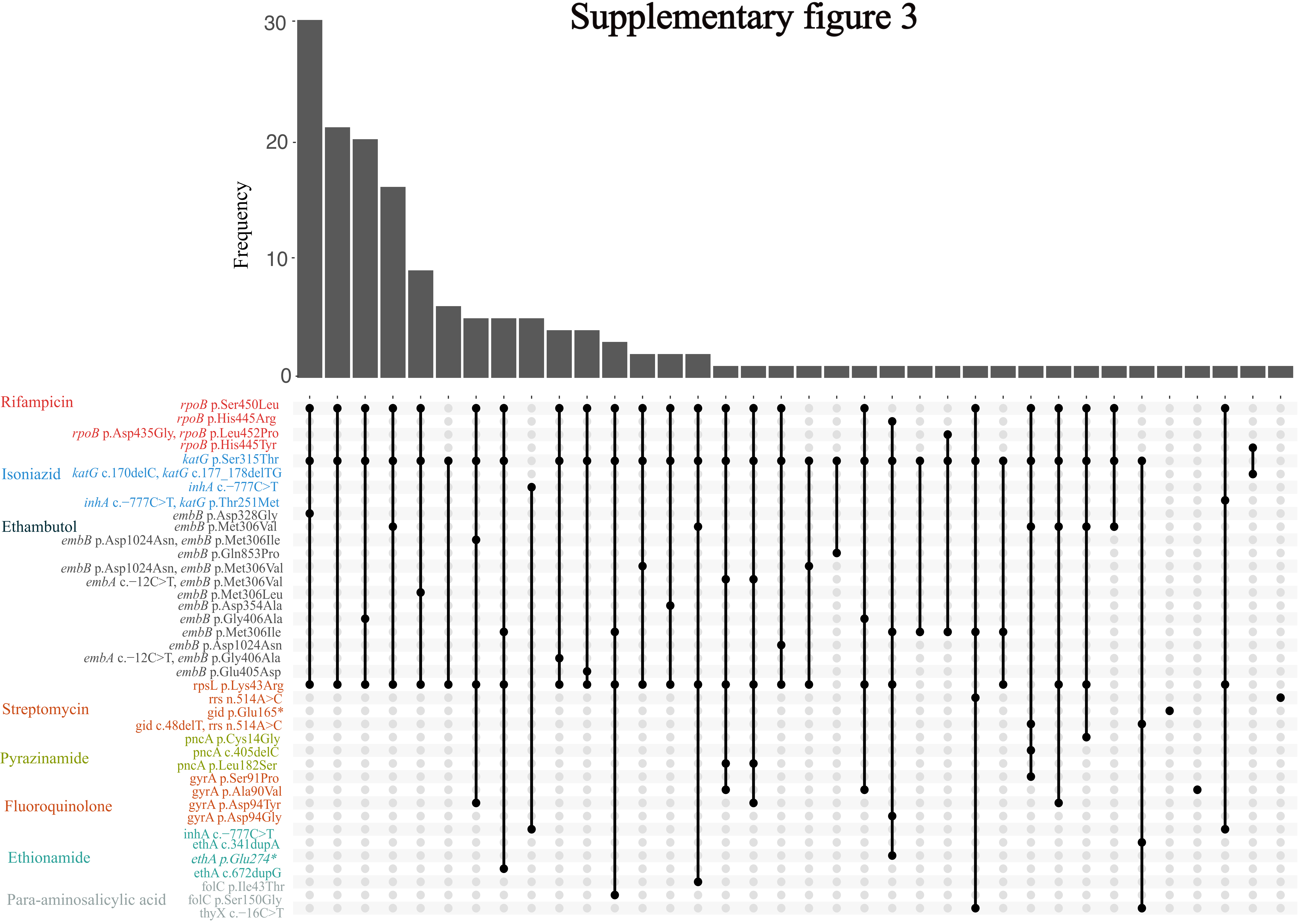

### Supplementary figure 4

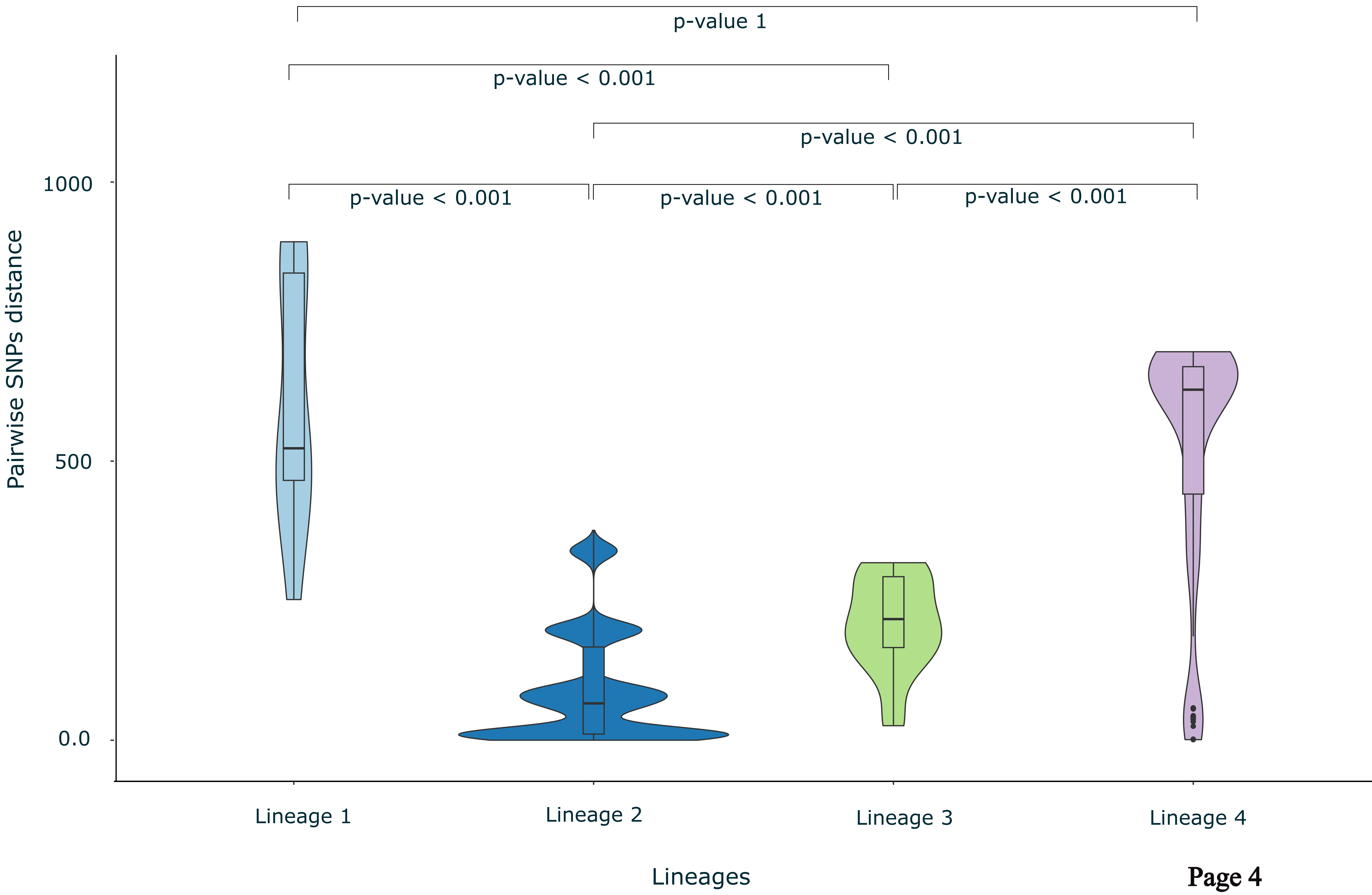

### Supplementary figure 5

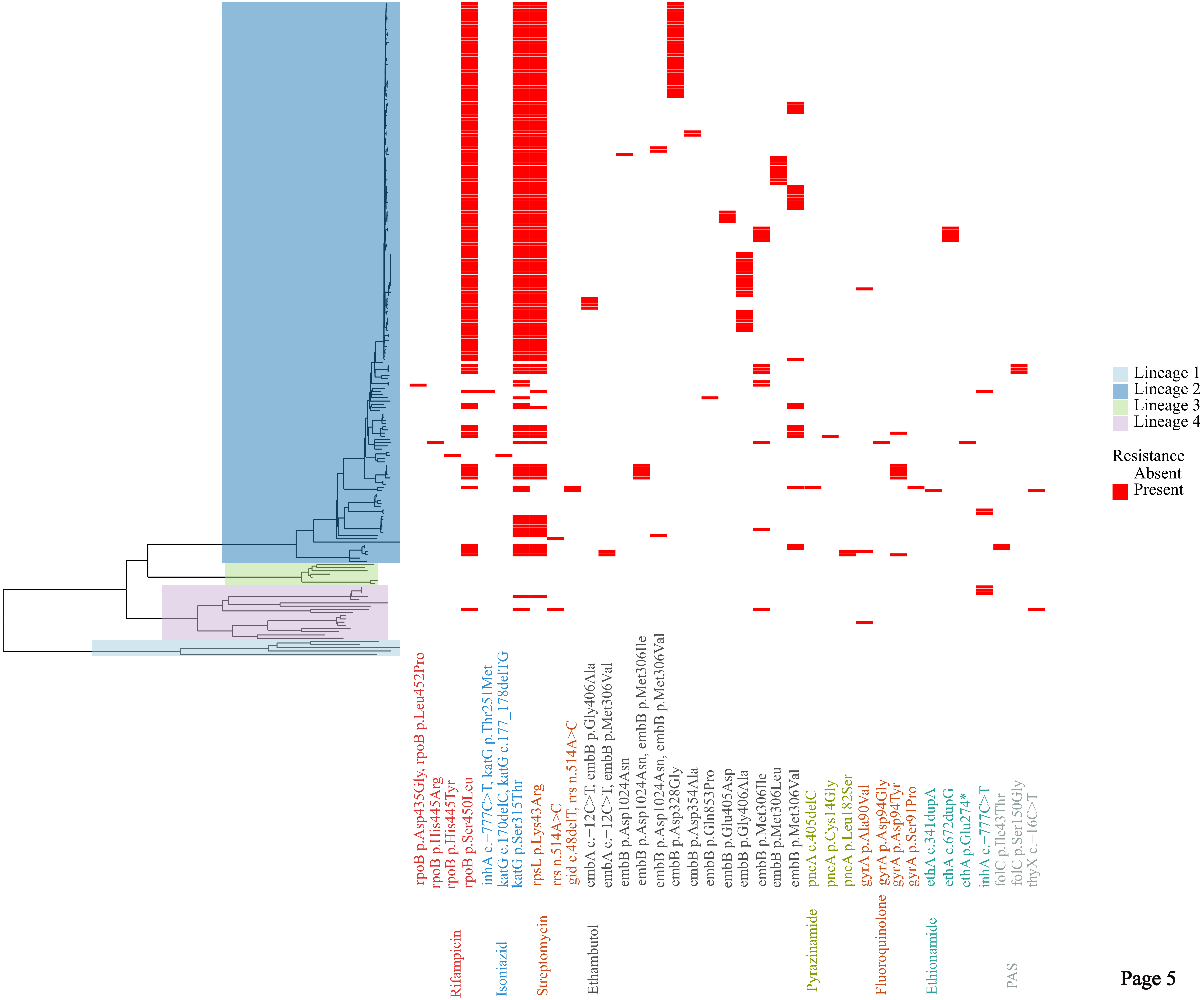

### Supplementary figure 6

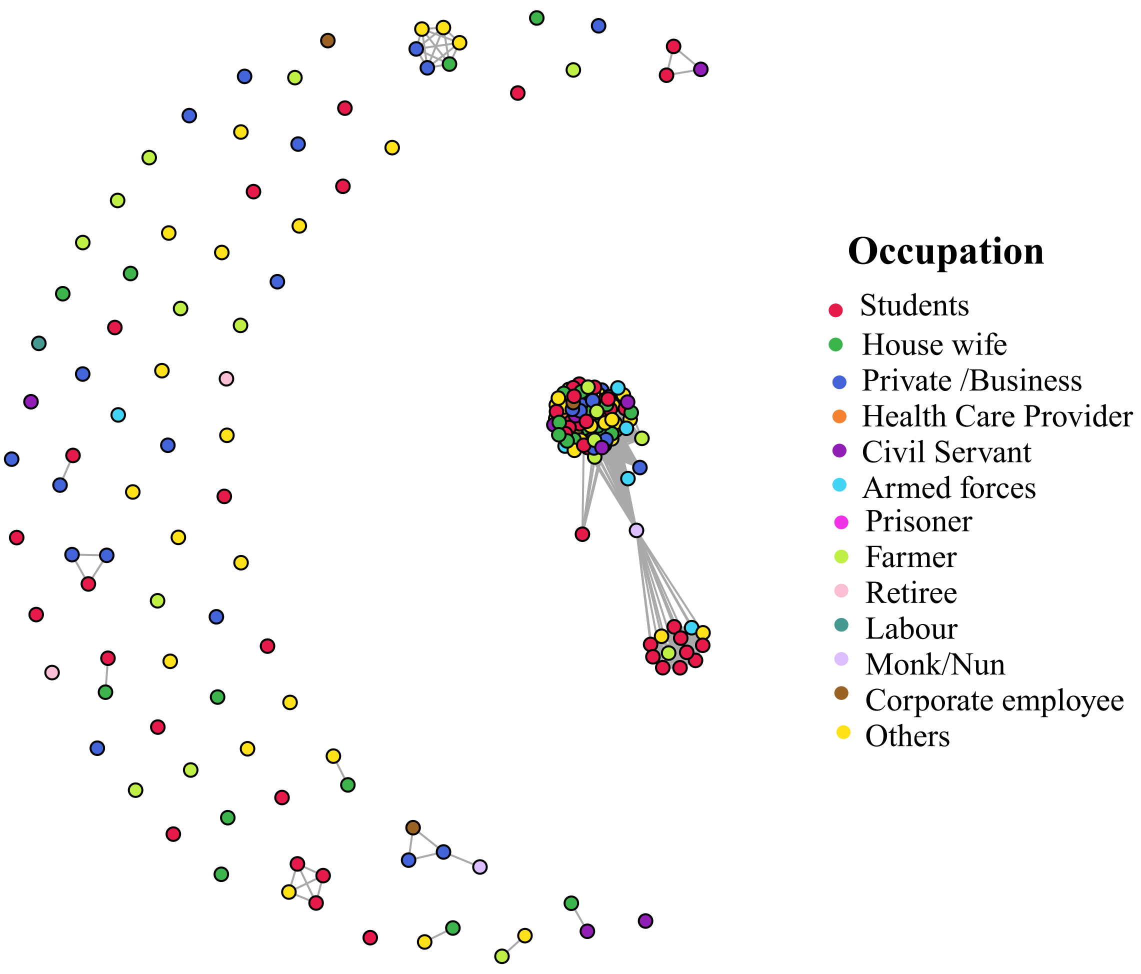

### Supplementary figure 7

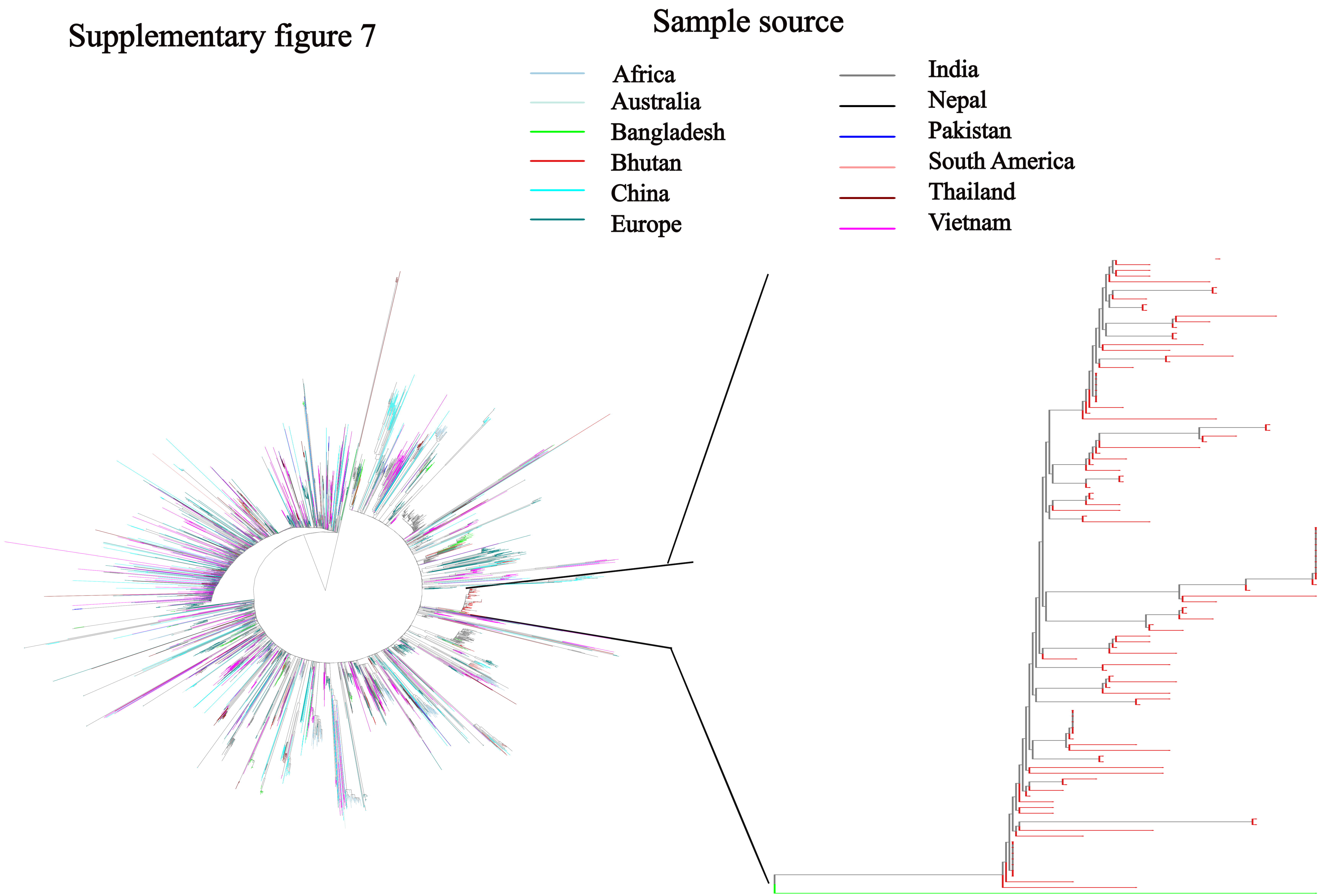
